# Effect of sample preprocessing and extraction methods on the physical and molecular profiles of extracellular vesicles

**DOI:** 10.1101/2023.07.17.549204

**Authors:** Lucile Alexandre, Molly L. Shen, Lorenna Oliveira Fernandes de Araujo, Johan Renault, Philippe DeCorwin-Martin, Rosalie Martel, Andy Ng, David Juncker

## Abstract

Extracellular vesicles (EVs) are nanometric lipid vesicles that shuttle cargo between cells. Their analysis could shed light on health and disease conditions, but EVs must first be preserved, extracted and often pre-concentrated. Here we firstly compare plasma preservation agents, and secondly, using both plasma and cell supernatant, four EV-extraction methods including (i) ultracentrifugation (UC), (ii) size exclusion chromatography (SEC), (iii) centrifugal filtration (LoDF), and (iv) accousto-sorting (AcS). We benchmarked them by characterizing integrity, size-distribution, concentration, purity and the expression profiles for nine proteins of EVs, as well as overall throughput, time-to-result and cost. We found that the difference between EDTA and citrate anticoagulants vary with the extraction method. In our hands, ultracentrifugation produced a high yield of EVs with low contamination; SEC is low-cost, fast, and easy to implement, but the purity of EVs is lower; LoDF and AcS are both compatible with process automation, small volume requirement, and rapid processing times. When using plasma, the LoDF was susceptible to clogging and sample contamination, while the AcS featured high purity but a lower yield of extraction. Analysis of protein profiles suggest that extraction methods extract different sub-population of EVs. Our study highlights the strength and weakness of sample preprocessing methods, and the variability in concentration, purity, and EV expression profiles of the extracted EVs. Pre-analytical parameters such as collection or pre-processing protocols must be considered as part of the entire process in order to address EV diversity and their use as clinically actionable indicators.

---

Extracellular vesicles (EVs) are nanometric lipid bilayer vesicles released by cells. They play a central role in cell-to-cell communication via ligand signaling and shuttling of cargo between cells including proteins, lipids and nucleic acids between the cells^1^. EVs are considered as highly promising biomarkers, notably for cancer^2^, as they can be directly collected by liquid biopsy, without the need for invasive procedures or surgeries. EVs are commonly characterized and identified by the expression of proteins; the tetraspanins CD63, CD9 and CD81 are ‘canonical’ EV markers^3^. The expression of these proteins is analyzed using a variety of methods, including western blots, mass spectrometry, flow cytometry, microarray, etc… Previous publications have shown that detailed protein analysis of EVs can reveal essential information on the evolution of a disease such as the development of a tumor^4^, Alzheimer’s disease propagation ^5^ or the spread of Parkinson’s disease^6^.

Prior to analysis, EVs must be collected, extracted and preconcentrated from the sample and the complex matrix. Blood, and by extension plasma and serum, is the most widely used biofluid for analysis and is readily collected via a phlebotomy. Indeed, the complex matrix of plasma can interfere with the analysis of EVs, and produce false positive signals. This puts stringent demands on the extraction methods which must content with high concentrations of potential contaminants. Plasma has been found to be a rich source of EVs and the number of EVs can be of the order of 10^10^-10^12^ particles per milliliter. While it is a large number, it corresponds to picomolar concentrations, considered vas ery low for proteins or metabolite. Hence, the preservation of EVs, as well as their purification and enrichment constitute critical pre-analytical steps to subsequent EV analysis.

Centrifugation remains the gold standard thanks to the high purity achieved by several steps^7,8^ and sometimes using gradient methods^9^. However, it is time-consuming, require expensive instruments and consumables. Size-exclusion chromatography is now widely used as a simpler, less expensive, and faster option^10,11^. Several other techniques have already proven their effectiveness, such as immuno-affinity^12,13^ or polymeric precipitation^14^. Microfluidics offers the option to work with smaller volumes, reduce costs, and automatize processes. New solutions have been developed on-chip, based on filtration^15^, acoustic^16^, viscosity^17^ or thermophoresis^18^. The need for simplified extraction is served by instruments that automate the workflow, such as the centrifugal dual-filtration system called Exodisc^19^ or the microbead-assisted acoustic trapping of EVs named AcouTrap^20,21^.

The heterogeneity of EVs constitutes a significant challenge to preprocessing and analytical procedures. EVs from different origins and types cover several orders of magnitudes in volume with size ranging from 50 nm to ~ 500 nm^22^. The membrane composition and cargo of EVs, either on their surface or encapsulated inside the lipid bilayer, is also heterogeneous. There is an effort to establish protein expression and identify subpopulations, however variations between studies and publications are apparent, and are often ascribed to lack of standardization of preprocessing and extraction steps. Whereas for membrane protein-based enrichment, there is a voluntary bias towards a particular subpopulation, other methods also enrich subpopulations of EVs based on physical (e.g., size), chemical (e.g., phospholipid composition) or molecular features. While not well understood, it is now widely recognized that each extraction method is biased in some way based on the molecular content or the size of the EVs, and as a result may fail to enrich certain subpopulations that then remain undetectable.

The standardization of EVs concentration, extraction and analysis remains a significant challenge. It limits our ability to compare results from different studies and to support the understanding of EVs and their biological function and clinical potential. The EV community promotes the need for standardization and multiple papers have been published, notably by the International Society of Extracellular Vesicles (ISEV)^23^ leading to the creation of new reporting technologies such as Exocarta^24^ or EV-track^25^. Hence, the performance of various EV concentration and extraction methods and their biases have been the subject of comparative studies^26,27^, especially the EV extraction from plasma^28^. However, in the absence of a ground truth with regards to what the pristine EV composition is, characterizing the absolute performance is not possible. Therefore, comparative methods are primarily used, and new methods benchmarked to existing ones to understand their performance and biases.

In this manuscript, using cell supernatant and plasma as samples, we evaluate four label-free EV enrichment methods: ultracentrifugation (UC), size exclusion chromatography (SEC) columns, and two emerging microfluidic technologies: centrifugal filtration on a Lab-on-a-disc (LoDF)^19^ and acoustic enrichment (AcS)^20^. EVs extracted from cell supernatant human squamous carcinoma cell line (A431) and pooled plasma samples were collected. Next, the extraction and enrichment of spiked-in and native EVs was characterized for three replicates. We also report the effect of blood collection tubes and pre-processing protocols. EVs in pre- and post-enrichment samples were characterized by Transmission Electron Microscopy (TEM), Tunable Resistive Pulse Sensing (TRPS), Nanoparticle Tracking Analysis (NTA) and antibody microarrays. The enrichment methods are benchmarked in terms of EV integrity, size-distribution, concentration, sample throughput, time-to-result, ease-of-use, purity, cost, as well as by comparing the expression profile of eight membrane proteins. Our study highlights the impact of pre-analytical steps and enrichment biases in terms of EVs sub-populations, contamination and changes in protein expression.

## Materials and methods

### EVs from cell supernatant

Human epidermoid carcinoma cell line A431 with CD63-GFP transgene^29^ was maintained in Dulbecco’s Modified Eagle Medium (Gilbco, USA) supplemented with 10% FBS and 1% PS (Gilbco, USA) under ATCC’s recommendation. To initiate EV collection, the normal growth media was replaced with DMEM media containing 5% EV-depleted FBS and 1% PS. When 80% confluency was obtained, the cell supernatant was collected, centrifuged at 400 x g for 15 min, and filtered using 0.22 um membranes (Filter Millex®?0.22 um #SLGPR33RS). Cell supernatant was concentrated to desired volume via centrifugation. For SEC and AcS extraction, this step was repeated.

### Plasma collection

Plasma was collected from healthy patients with informed consent form. The collection was approved by the McGill research ethics office (IRB study #A05-M27-16B). Blood samples were collected by a certified nurse and anonymized. The blood was collected in EDTA (VWR, CABD367861L) and citrate (VWR, CABD369714L) tubes, stored at 4°C, and processed within 4h of collection. The collected blood was then centrifuged (Thermo Fischer Scientific, Sorvall RT1 centrifuge) 15 min at 1100 x g then frozen at - 80°C. For extraction assessment, plasma samples were spiked with A431 EVs at the adjusted concentration of 10^10^ part/mL.

### Pre-analytical parameters

Three different sample pre-processing methods are compared in this article: first, samples were thawed at room temperature before use, no additional centrifugation or filtration step; the second method relies on a second step of centrifugation at 1100 x g for 15 min before storage at -80°C. After storage, samples were thawed at room temperature, centrifuged at 2500 x g for 15 min and filtered through 0.22 μm membrane; finally, samples were thawed at room temperature, centrifuged first at 4700 g for 15 min then at 14000 g for 30 min and finally filtered through 0.22 μm membrane. Collection of samples was performed under the ethic approval of the study A04-M36-13B (IRB number).

### Extraction methods

Ultracentrifugation extraction was performed by pouring a 1-2 mL sample in a polycarbonate tube and diluted in PBS up to 8 mL. The diluted sample was then centrifuged using a Beckman ultracentrifuge (Optima L90K) with two cycles of 1h10 at 110,000 x g at 4°C (rotor type 70Ti, Beckman Floor Ultracentrifuge Optima L90K). Between them, the supernatant was discarded and the pellet was resuspended in 8 mL of PBS. The final pellet was resuspended in 150 μL of PBS. SEC was performed using qEV SEC columns, purchased from Izon® (qEVoriginal/70nm/500μL), following their commercial protocol. 500 μL preconcentrated samples were flown through the columns and only fraction 8 was collected as eluate. The acoustic extraction was performed with the AcouSort system developed by AcouSort®. 50 μL of a suspension of 1 μm diameter beads were injected through the system at 50 μL/min followed by 90 μL of preconcentrated cell supernatant or 50 μL of plasma diluted 1:2 at 15 μL/min and by 200 μL of PBS at 25 μL/min. The extracted EVs were released in 50 μL of PBS at 1 mL/min. The process was repeated to match the volume of sample to analyze. A last step of centrifugation at 500 x g for 2 min was performed to separate the sample from the beads. Finally, the LoDF system was purchased from LabSpinner®, Exodisc C and P. The disc filters were first wetted by 1 mL of PBS, then 2 mL of cell supernatant or 100 μL of plasma sample was flown through the system followed by a washing step of 100 μL of PBS. The 100 μL supernatant block in between the filters is then collected, the chamber is washed with 100 μL of PBS. Volumes of the samples analyzed are summarized in Table S1.

### Characterization methods

EVs characterization included NTA (NanoSightNS500), TRPS (Izon®), Nanodrop (spectrometerND1000), TEM (FEI Tecnai 12 120 kV) and a custom antibody microarray for protein profiling^30^ (description in supplementary).

### Analysis

All results are plots with each data point, means and standard deviation (SD). Drawing and analysis was performed using GraphPad Prism. For plasma samples, the normalized final concentration of extraction is the final concentration after extraction related to initial and final volumes of samples (eq1 in Supplementary). For the heatmaps, the fluorescent signal corresponding to a protein (capture on the slide) is normalized by the signal corresponding to the same capture spot recorded in buffer. For each graphic, clustered analysis is run to organize columns order on Matlab (function *clustergram*), based on Euclidian distance, with the drawing of the corresponding dendrogram, then the signal is normalized by the highest measured signal to obtain a percentage value.

## Results

### I. EV-extraction methods

We evaluated four EV extraction methods using cell supernatant and subsequently plasma samples, including (i) UC (ii) SEC (iii) LoDF and (iv) AcS. The protocols are detailed in the methods and illustrated in Fig. 1. Briefly, UC was conducted using a standard protocol including two cycles of centrifugation 110,000 x g at 4°C for 70 min each, with dilution in order to minimize contamination. SEC was conducted using Izon® columns according to manufacturer’s protocol and fraction 8 was collected for further analysis. AcS uses the so-called AcouTrap™ for trapping 1-μm-diameter beads that produce acoustic streaming and vortexes that in turn trap EVs; EVs are too small for direct acoustic trapping. LoDF was using centrifugal flow across a spinning disk called Exodisc™ and flowing the sample through a first filter with 600 nm pores that retains large particulates, followed by a second filter of 20 nm pores that retains EVs.

**Figure 1.**
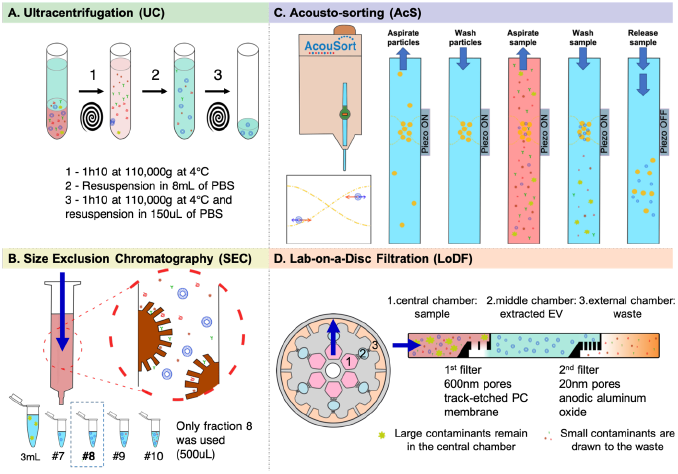
Description of the methods of extraction of EVs: **(A)** UC extraction was performed by two successive cycles of centrifugation and dilution of the sample, the final pellet was resuspended in 150 μL of PBS **(B)** SEC was performed on 500 μL samples following the manufacturer’s protocol. Final volume was 500 μL **(C)** For AcS samples, EVs were trapped in between 1 μm diameter beads. The extracted EVs were released in 50 μL of PBS **(D)** The LoDF system is composed of two filters with pores of 600 and 20 nm. Extracted EVs were collected in-between filters for a final sample of 200 μL.

The characterization was performed starting with cell supernatant samples to limit the number of contaminants and extended to human plasma which has a complex matrix and is clinically more relevant.

### II. Characterization of EVs extracted from cell supernatant

#### a. Comparison of size and concentration of extracted EVs

EV supernatant samples from A431 cell line were collected and processed with one of the four extraction methods on the same day. The total protein concentration was measured by UV absorption at 280 nm, Figure 2A. The concentration of proteins was comparable for UC, SEC, and AcS, while LoDF samples had much higher UV adsorption as a result of higher protein concentration. Next, the EV concentration and size distribution was measured by TRPS^31^. The particle concentration for LoDF was about tenfold higher compared to UC and SEC, while AcS yielded an overall lower concentration of particles along with larger variability, Figure 2B. For LoDF the average size of the extracted particles was ~ 15% smaller, Figure 2C & D. The smaller average size could be related to better yield in extracting smaller EVs, or alternatively, reflect contamination by other particles possibly comprising proteins, nucleic acids, lipoproteins and mixtures. To evaluate which one of the two scenarios is applicable, we analyzed the samples by TEM (Supplementary Figure S1). We observed a higher number of contaminants with diameters under 30 nm for LoDF consistent with higher particle counts and smaller average diameter. These results are consistent with the literature on Exodisc® LoDF ^26,32^.

**Figure 2.**
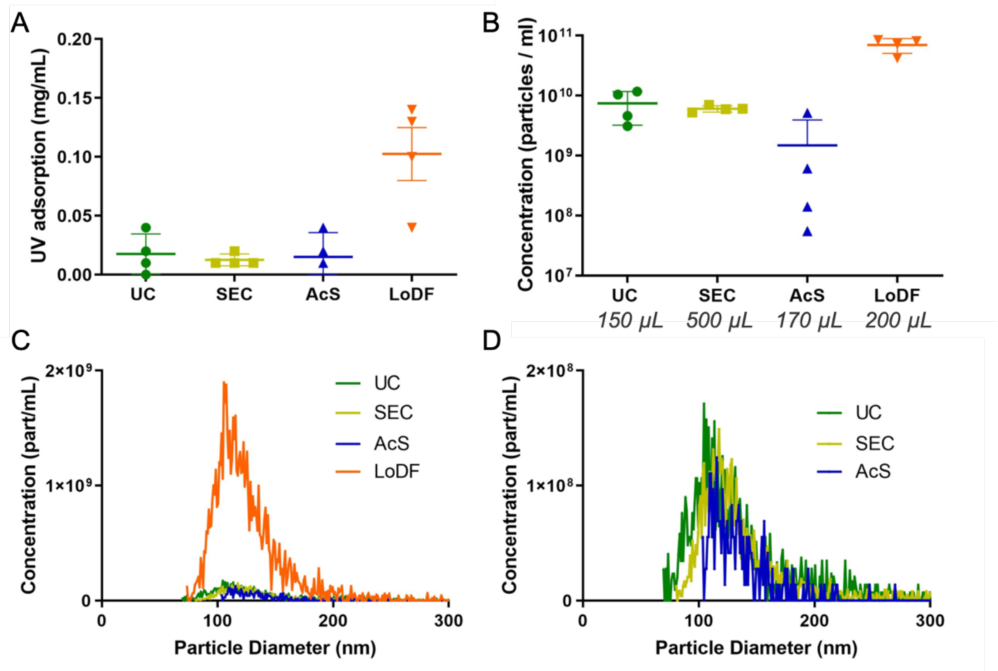
Characterisation of (A) protein content, (B) final concentration and volumes of samples after extraction and **(C-D)** size distribution of EVs extracted from cell supernatant using UC, SEC, AcS and LoDF by TRPS. Samples processed using the LoDF have the highest protein content, and the highest concentration of EVs while EV size is smaller than with the three other methods. AcS samples had a lower concentration of particles along with larger variability (4 replicates showing mean and SD as error bars)

#### b. Characterization of the protein expression of extracted EVs

The protein expression profile of extracted EVs was evaluated by an in-house microarray immunoassay (Figure 3 A & B). The microarray with antibodies against EV surface proteins was incubated with sample containing 1.5 10^9^ particles per mL, and bound EVs incubated with a cocktail of biotinylated antibodies against the three tetraspanins CD63, CD9 and CD81, followed by fluorescent-labelled Streptavidin (Figure 3 C). Transmembrane and cytosolic proteins, as advised by the MISEV2018 guidelines^23^, were included in the panel of capture antibodies to help confirm the integrity and nature of EVs.

**Figure 3.**
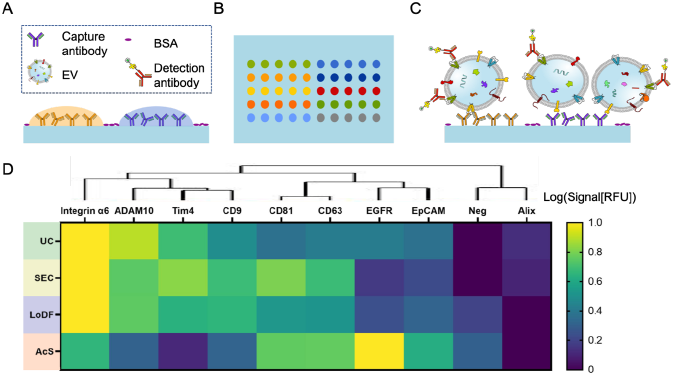
Antibody microarray-based profiling of protein expression of EVs extracted from A431 cell supernatant. **(A)** Schematic of antibody spots, (**B**) of array layout, and (**C**) of immobilized EVs with fluorescently labelled detection antibody cocktail against CD9, CD63, CD81. (**D**) Heatmap of protein expression in EVs. Tetraspanins and integrins are generally highly expressed, while the expression levels can vary according to the extraction method. The low expression of Alix indicates that EV integrity was preserved. The binding of EVs to TIM4 was observed for UC, SEC and LoDF, as expected, but not AcS. Conversely, EGFR expression was low everywhere except in AcS EVs, suggesting that AcS extracts a markedly different subpopulation compared to other methods. These results underline that method will enrich for different sub-populations of EVs.

Assuming that the expression of tetraspanins is conserved across the four methods, and neglecting the influence of changes in average EV size, then the relative fluorescent signals for each method will reflect the relative enrichment or depletion of EVs immobilized by the capture antibody. The lower expression of those proteins in LoDF samples, Figure S2, would be consistent with higher levels of contamination, or a bias towards smaller EVs that may express lower number of proteins. The integrin α6, involved in cellular adhesion^33^, is highly expressed by EVs from A431 cell line. The epidermal growth factor receptor EGFR and the metalloproteinase ADAM 10 were selected based on the reported high expression in this cell line^34^. EpCAM is also a recommended marker for EVs and known to be overexpressed in A431 cells. Interestingly, we observed significant differences in expression of these proteins in the EVs as function of the extraction method. TIM4 (T cell immunoglobulin and mucin domain containing 4) binds phosphatidylserine that is exposed on many EVs ^35^, and was previously used for EV isolation ^36,37^. Here we used it to assess the binding of EVs independently of proteins expression. The low phosphatidylserine expression in EVs extracted by AcS combined with difference in expression of other proteins suggest that AcS may be extracting different sub-populations of EVs. The integrity of EVs was confirmed by the absence of signals for the cytosolic protein Alix (ALG-2-interacting protein X).

### III. Characterization of EVs extracted from human plasma samples

We evaluated the performance of the four extraction methods to enrich EVs from plasma collected from healthy patients in EDTA and citrate tubes spiked with previously analyzed A431 EVs, using the processing protocol established above.

#### a. Characterizing sample pre-processing on EVs collection

To better understand the impact of pre-analytical variables, we compared three sample pre-processing (SPP) protocols, from simpler to more complex

SPP1: Collect plasma → freeze → thaw

SPP2: Collect plasma → centrifuge (1100 g) → freeze → thaw → centrifuge at low g (2500 x g) → filter through 0.22 μm membrane

SPP3: Collect plasma → freeze → thaw → centrifuge at moderate g (4700 g) → centrifuge at high g (14000 x g) → filter through 0.22 μm membrane

Figure 4 shows protein content measured by UV adsorption, number of nanoparticles, and average nanoparticle size for SPP1 – SPP3 using either EDTA or citrate tubes for blood collection and plasma separation. A general trend observed is that the more steps the protocol includes, the more loss of EVs is observed, while purity increases, as one might expect. UV 4adsorption measures EV associated proteins and free proteins, and reflects this trend. Again, LoDF yields ~ 10 times higher protein concentration likely due to contaminants, while AcS show low protein concentration consistent with low yield but high purity, both replicating trends observed in cell supernatant extraction. TRPS measurements reveal a low particle concentration for AcS that falls below 10^9^ part/mL, which is below the threshold of particle numbers needed for some analytical tools and applications. The reduction in average particle size following additional centrifugation step for SPP2 and SPP3 reflect the removal contaminants such as cell debris as well as of EVs.

**Figure 4.**
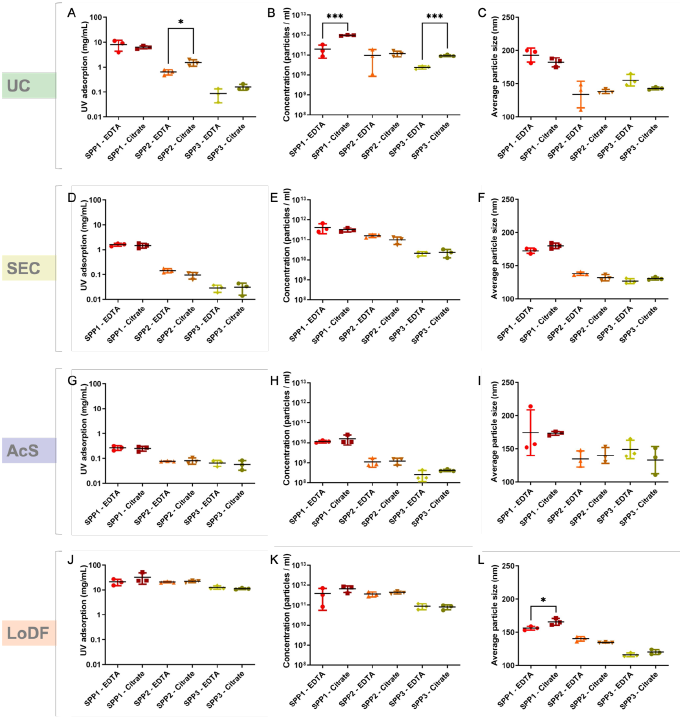
Characterization of the effect of sample collection tubes, pre-processing and extraction method on EV protein content, concentration and size for plasma from health volunteers. Total protein content as measured by UV absorption at 280 nm **(left column, A, D, G and J)** on nanodrop, concentration **(middle column, B, E, H and K)** and average size (**right column. C, F, I and L)** of the particles measured by TRPS. The EVs were extracted from both EDTA and Citrate tubes samples in three conditions: thawed, unprocessed samples (SPP1), additional centrifugation before storage, thawing, low g centrifugation (SPP2), additional moderate g centrifugation following thawing (SPP3). SPP3 yielded the highest purity. (mean shown as black bar, data and SD in in color)

The overall protein concentration, particle number and average diameter are not significantly different whether Citrate or EDTA were used as anticoagulant, except for UC extraction. EDTA samples seem more variable among the replicates whereas citrate samples were consistent, especially for UC, however more experiments are needed to confirm this trend.

Based on these results, and because contamination of EVs samples by non-EV proteins and particles is a major concern for EV analysis, the SPP3 protocol was selected for subsequent experiments as it yields purer EV samples.

#### b. TEM imaging of EVs

To evaluate the integrity of plasma EVs following SPP3 and extraction, we imaged them by TEM, Figure S3. For AcS samples with low EV concentration, a larger volume was used to obtain a comparable number of EVs. Negative staining, using uranyl acetate, was used to enhance contrast between the particles and the background including contaminants. We identified round and intact EVs identified by their ‘cub-shape’^38^ of different size in samples obtained by every extraction (see white arrows); particles were smaller in LoDF samples.

TEM images can be used as a proxy for qualitatively assessing concentration but with the caveat that they constitute only a small sample that may not be representative, and that EVs may not be distributed homogeneously on the TEM grid. TEM images of LoDF samples revealed a higher concentration of particles relative to other samples, consistent with the literature^32^. For AcS samples, the concentration of EVs was too low, hence a larger sample volume was used. The images of AcS extracted EVs are comparable to particles observed in other publications^20^ where the same difference in size was also observed.

Contamination of samples can also be evaluated via the TEM images, Figure S3. Differences in the level of grey in the background of TEM images are related to the density of contaminants such as proteins, salt and aggregates. In the images of samples extracted by SEC columns, long filaments were observed which are known residues from sepharose resins^39^. Samples extracted by LoDF had a high concentration of contaminants, consistent with previous publications^32^. Round and bright particles that are distinct from ‘cup-shaped’ EVs were identified as lipoproteins. However, this affirmation was altered by the emergence of exomeres (i.e. EVs < 50 nm) which have a similar round and bright appearance than lipoproteins and do not adopt a cup-shape. There is now ambiguity as to whether the bright spots are contaminants or exomeres.

#### c. Comparison of protein concentration, EVs concentration and size distribution

We evaluated the total protein concentration, particles concentration and size distribution by UV absorption, see Supplementary Figure S4, TRPS and NTA measurement of plasma samples collected in citrate and EDTA tubes, processed according to SPP3 and extracted using each of the 4 methods. As plasma samples are more complex than cell supernatant ones, we measured the concentration and size by both TRPS and NTA. NTA tracks particle diffusion thus deriving the coefficient of diffusion and the particle diameter^40^. Coupling TRPS and NTA provides a more complete characterization of the EVs profile. A normalization process was necessary because unlike for cell supernatant, the starting sample volume of plasma for each method was not the same, Table S1. The final concentration was corrected by the ratio of final to initial volumes to produce a result that represents a yield of extraction. The final concentration of the extracted samples before and after normalization by volume, as explained in Materials and Methods, are shown.

UC produces a high concentration in extracted samples, while SEC offers a higher yield of extraction, after normalization. Differences in measured concentrations between NTA and TRPS are significant only for AcS samples, whose final concentrations are lower than the ones from other samples. The discrepancy could be explained by the differences in limits of detection of NTA and TRPS. LoDF extraction resulted in a high concentration of proteins, more than 50 times higher than other methods. TRPS and NTA measurements revealed a high concentration of nanoparticles after extraction for LoDF compared to UC, SEC and AcS, Figure 5A & B, that was preserved after normalization, Figure 5C & D. However, both for EDTA and citrate plasma, a high concentration of contaminants was observed by TEM which could both contribute to increase in protein concentration and nanoparticle count, and the broad size distribution. The distribution and ratio of contaminant nanoparticles vs EVs could not be established and will require further characterization.

**Figure 5.**
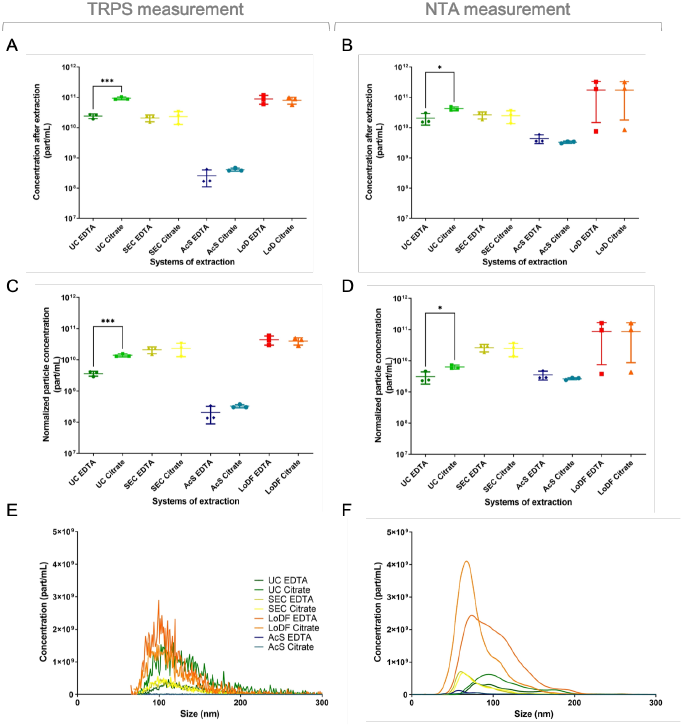
Characterization of the concentration, yield and size distribution of EVs extracted from EDTA and Citrate plasma samples using UC, SEC, AcS and LoDF by TRPS (A, C and E) and NTA (B,D and F). A high yield of extraction was measured for LoDF and SEC extraction. For all samples except AcS, the particle concentration as measured by TRPS and NTA are consistent; for AcS, TRPS yielded a lower concentration than NTA that can be related to a low final concentration compared to other samples. Measured size of the particles is higher when measured by TRPS, as observed on measured profile (E and F) between NTA and TRPS. Experiments included 3 replicates and with error bars representing SD and showing a line for mean.

Whether EDTA or citrate tubes were used did not affect the composition of extracted EVs for SEC AcS and LoDF samples in our hands. However, for UC, the protein concentration appeared higher (almost twice as high, although not meeting statistical significance) in citrate samples compared to EDTA ones, Figure S4. Concentration of EVs were also higher in UC Citrate samples compared to EDTA ones, Figure 5A & B.

No obvious difference in size of extracted EV was noticed in-between the two anticoagulants used for those experiments. Measured size distribution of the particles is shifted toward bigger sizes when measured by TRPS compared to NTA for all samples, Figures 5E & F, S5 & S6. It correlates with the fact that NTA detects higher number of small EVs than TRPS^41^. Our data suggest that UC allowed to extract populations of bigger EVs than SEC, AcS and LoDF, as it can also be observed on TEM imaging, Figure S3.

#### d. Protein expression characterization of extracted EVs

An antibody microarray was used to assess the multiplexed protein expression of EVs captured using a cocktail of Anti-CD9, -CD63 and -CD81 antibodies, Figure 6 (see materials and methods for details). The analysis for AcS EDTA and AcS Citrate samples was performed at a lower concentration. Plasma samples were spiked with cell supernatant A431 EVs analyzed previously, with the expectation of detecting both spiked and native EVs.

**Figure 6.**
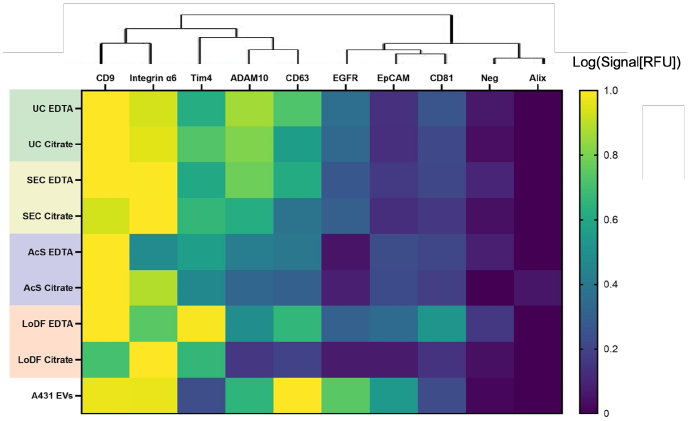
Protein expression in EVs from A431 cell supernatant spiked into plasma profiled by antibody microarray. A431 cell supernatant EVs were spiked into plasma then the sample were extracted by UC, SEC, AcS and LoDF from the EDTA and Citrate plasma tubes and incubated on the microarray. EVs were immobilized by capture antibodies (top labels), and then probed with a cocktail of fluorescently detection antibodies against CD9, CD63, CD81, and after rinsing, the fluorescence signal detected. Log values of the signals were normalized by the highest protein expression for each sample. For UC, SEC and LoDF samples were adjusted to 10^10^ part/mL and AcS 10^8^ part/mL. As expected CD9 was highly expressed in all plasma samples, and differences in proteins expression are less visible due to the high number of EVs in plasma. The protein profile of EVs from A431 cell supernatant prior to spiking into plasma is shown at the top as a guide. Note that both spiked EVs and native EVs are detected in the plasma samples.

As explained, the panel of protein targets included transmembrane proteins, phosphatidylserine (recognized by TIM4) and cytosolic proteins serving as negative control. Tetraspanins and integrins are highly expressed in all plasma samples. However, it is interesting to notice that clustering may differ between cell supernatant and plasma sample. CD9, ADAM 10, Integrin α6 and Tim4 are highly expressed in all samples, whereas no signal was measured for Alix. However, CD63 and CD81 are clustered in cell supernatant samples but not in plasma samples. EVs extracted from plasma samples expressed highly CD9^42^, compared to EVs extracted from cell supernatant. We conclude that the signal from native EVs dominated the one from spikedin EVs form this cancer cell line. EpCAM and EGFR expression, specific to the A431 cell line was decreased on the other hand, which along with other EVs was subdued by the signal from the plasma sample. As expected, a low signal was recorded from LoDF samples and ACS samples, related to high contamination and lower concentration, respectively, as well as lower consistency (See Figure S7). When using EDTA compared to citrate, UC and SEC had a slightly higher protein expression, while AcS and LoDF had a lower one.

## Discussion

Previous publications have evaluated new technologies of extraction^19,20^, extraction from different media^32^, or compared preanalytical parameters for EV collection and storage^43,44^. We provide here a comparative study that evaluates the sample process flow, sample nature and composition, extraction protocols and methods, to identify important parameters separately as well within a process flow and identifying co-dependencies and optimal process flows. We compared collection protocols and four methods of extraction: (i) UC, (ii) SEC, (iii) AcS and (iv) LoDF for isolation of EVs from cell supernatant and human plasma. We optimized pre-analytical parameters for plasma pre-processing, such as collection tubes and various centrifugation steps, to match usual hospitals protocols and purity requirements, and finding that two steps of centrifugation after thaw and a 0.22 μm filtration allow to reduce contamination. Finally, we evaluated the impact on the EV protein profile based on our home-built immunoassay.

Our study corroborated the high purity of EV samples obtained by UC, while also facing lengthy process time of this method, and difficulty to gain access in our case. To qualitatively compare the strength and weakness of each of the four purification methods for cell supernatant and plasma, we benchmarked their performance according to five and six criteria, respectively including (i) final concentration, (ii) speed, (iii) volume preconcentration factor, (iv) purity, (v) accessibility, and in the case of plasma, (vi) yield, Figure 7. The performance of different methods for processing cell supernatant showed a significant variation for most parameters, Figure 7A. No individual method outperforms the others, and each method has advantages (high purity for UC, accessible for SEC, fast for AcS and high yield for LoDF) and drawbacks (lengthy for UC, low preconcentration for SEC, low final concentration for AcS and high contamination for LoDF). The analysis of the protein expression in the purified EVs revealed differences between all methods, suggesting bias and selective enrichment or depletion of subpopulations of EVs. Bias introduced by the extraction method means that comparative studies across sites need to account for it, and harmonize extraction protocols and methods, develop strategies to benchmark and compensate for possible bias.

**Figure 7.**
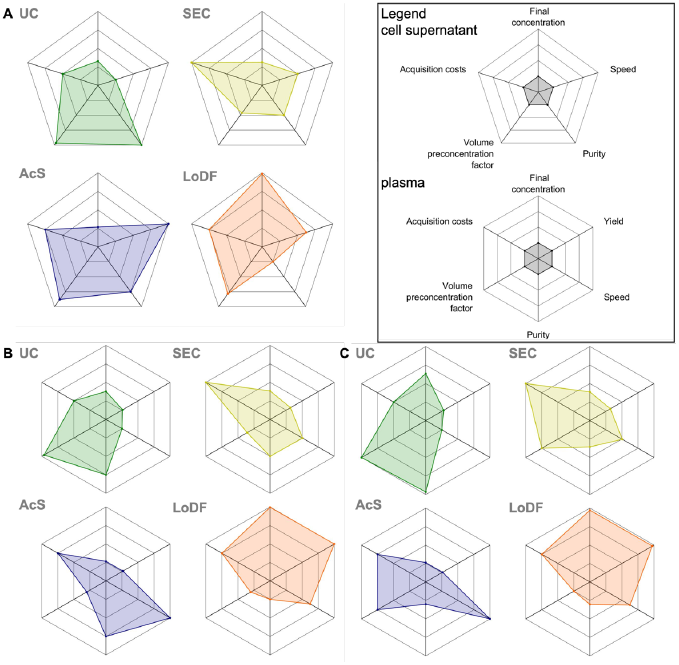
Evaluation of the four methods of extraction: UC, SEC, AcS and LoDF for **(A)** cell supernatant samples in term of speed, volume preconcentration factor (ratio of the final to initial volume), recovery rate, purity and accessibility and **(B)** EDTA and **(C)** Citrate plasma according to the same criteria plus yield. UC shows excellent performance in purity and preconcentration, but accessibility, yield, speed, and final concentration are not optimal. SEC, AcS and LoDF have interesting characteristics that make them challenging. However, all radar charts are unbalanced, showing a remaining need for simple, automated EV extraction method.

A pre-analytical protocol is necessary prior to extraction of EVs from plasma and three different ones were tested, Figure 3, and their influence on the purity and profile of the extracted EVs characterized for each extraction method, and the best combinations identified. The method of extraction also impacts the properties of the extracted EVs, Figure 7B & C. UC provide high volume preconcentration, as the pellet from a large sample can be resuspended in a volume of buffer as low as 150 µL, inducing a high final concentration with exceptional purity. However, it is a lengthy process, the yield is low considering the high initial volume, and accessibility can be a challenge if an ultracentrifuge is not available. SEC is an accessible alternative with mid-range performance on many criteria. The final concentration is lower than with UC but with the higher yield of extraction. SEC is interesting for studies where the volume of samples is small (from 500 µL to 2 mL), and is most easily implemented as it does not require additional equipment. AcS offers a fast extraction with high purity, and if the equipment is available, with low-cost consumables, easy to integrate in a laboratory and to operate, but the final concentration, yield and preconcentration factor were found to be comparatively low. It is worth mentioning that the yield of extraction is close to the one from UC. LoDF also depends on a specialized, compact equipment along with proprietary consumables, and produces high yields of proteins and nanoparticles, with reasonable time-to-result, and an easy-to-use range speed and easy-to-use technology, but suffers from a low volume preconcentration factor and a comparatively high contamination. The choice of EDTA or Citrate tubes for blood collection overall had little impact on the performance of the various enrichment methods, except for UC, and possibly for SEC. Indeed, citrate plasma compared to EDTA plasma yielded higher final concentration of EVs with UC, see Figure 5, while purity of EVs extracted by SEC was higher when judged based on TEM images, see Figure S3.

This study further documented how the method of extraction could change the measured protein profile of the extracted EVs, suggesting that specific sub-populations of EVs are preferentially enriched. Lacking a standard and a ‘ground truth’, it is not possible to identify whether a method introduces a bias for any given protein, and comparisons could only be made relative to one another, and hence were not included in Figure 7. Interestingly, the subpopulation (by protein expression) of EVs enriched depended on the extraction protocol for both sample types (cell supernatant or plasma), as revealed by the different hierarchical self-clustering based on protein expression (compare Figure 3 and Figure 6). High expression of proteins, such as tetraspanins and integrins, can hide disparity on the expression of rarer proteins. Quantifying protein expression and coexpression beyond the canonical tetraspanins is important to consider when evaluating a processing pipeline as it can enrich or deplete certain subpopulations. Differences in protein expression and subpopulation enrichment were observed for cell supernatant based on the antibody microarray heatmap profiles. For plasma samples, the heatmap profiles showed less variation. Note that the microarray measured both spiked EVs and native EVs in the plasma that could not be distinguished, see Figure 6. Hence, the high CD63 expression of the spiked-in cell supernatant was not reflected in the measurements in plasma that showed high CD9, and in many cases high integrin α6, suggesting that native EVs were more abundant or possible matrix effects. Matrix effects could also reduce overall sensitivity and account for other differences. The effect of EDTA or Citrate collection tubes did also lead to heatmap intensity changes in protein expression profiles, but visually the overall pattern across methods of extraction was preserved, except for LoDF. When using citrate tubes, the expression of CD9 was markedly reduced compared to all other measurements. Further experiments will be needed to confirm these results, and to identify the mechanisms that may lead to differences in the abundance of EV subpopulations.

A limitation to our study is that we did not consider or evaluate the impact of sample storage and of different storage conditions. The four methods of extraction were performed on the same day, following the same initial sample towards minimizing any influence of storage, which will need to be studied given that many samples are stored prior to analysis. A better understanding and comparison of the available methods of extraction will help standardization of protocols, a need that is also reflected in analysis of the impact of storage conditions on EVs populations ^43,44^.

## Conclusion

This study demonstrates the importance of pre-analytical parameters in EVs studies, when working with either cell supernatant or plasma samples, and their impact on sub-sequential analysis, with a focus on protein expression analysis. We worked with multiple pipelines for sample collection and preparation, designed to match usual protocols from hospitals and labs, combined with four systems of extraction. We provide here new data on comparison of different combinations of collection and extraction parameters, showing that pre-analytical parameters directly influence the studied EVs.

We introduce here new keys for characterization and comparison of the pre-analytical parameters, to help scientists build their own pipeline of EVs extraction, while been aware of the compromises that need to be made and their impact on sub-sequent analysis. We believe that this step is essential to understand the bias introduced by the preparation of samples and it should be part of any EV analysis. The optimal sample collection protocol may not be a general characteristic, but one that depends on the purification method, and of course on the intended use and measurement method. Ignoring possible effects of sample processing could lead to inconsistent results even when only apparently minor changes are made, and could possibly lead to irreproducible or contradictory results. Pre-analytical parameters must be considered as part of the entire process, including which extraction method and which type of EV subpopulation is to be considered.

## ASSOCIATED CONTENT

### Supporting Information

Supporting information included a document presenting detailed Materials and Methods and supplementary figures (PDF)

## AUTHOR INFORMATION

## Author Contributions

The manuscript was written through contributions of all authors, all authors have given approval to the final version of the manuscript.

## ACKNOWLEDGMENT

This project has received funding from the European Union’s Horizon 2020 research and innovation programme under the Marie Skodowska-Curie grant agreement No 896313.

## Notes

### Competing Interest Statement

The authors have declared no competing interest.

