## Supplementary for "Effect of sample preprocessing and extraction methods on the physical and molecular profiles of extracellular vesicles"

### Materials and Methods:

*Chemicals:* All solutions were prepared with deionized water from a MilliQ system (resistivity of 18 MΩ cm, Millipore, Burlington, MA). Phosphate buffered saline (PBS, 10X, ph=7.4 [-] Calcium Chloride [-] Magnesium Chloride #70011-044) was purchased from Gibco and diluted to 1X concentration in MilliQ water. Bovine serum albumin (BSA) was purchased from Jackson ImmunoResearch laboratories (#001-000-162 Bovine Serum Albumin IgG-Free, Protease-Free). Tween20 were purchased from Sigma-Aldrich (BioXtra #P7949). Antibodies were purchased from R&D Systems, Biolegend, Sigma, Invitrogen and Life technologies. Streptavidin-AlexaFLuor 647 was purchased from Sigma Aldrich (#511223).

*Nanoparticle Tracking Analysis (NTA, NanoSightNS500):* EV samples were diluted in PBS to achieve a concentration in between $10^8$ and $10^9$ particles/mL. Samples were loaded in a syringe pump connected to the flow cell chamber. For each sample, 3 measurements were taken and averaged to calculate the particle concentration.

*Tunable Resistive Pulse Sensing (TRPS, Izon®):* The instrument was primed with silica particles of 100nm diameter, using a pore size membrane NP150. EV samples were diluted in electrolyte buffer then loaded onto the top fluid cell. The signal was recorded consecutively for a maximum of 10 minutes.

*Nanodrop (spectrometerND1000):* Measurements were performed using the A280 mode following the protocol of the supplier.

*Transmission Electron Microscopy (TEM, FEI Tecnai 12 120 kV)*: After extraction, EVs were store at -4°C. Then 5 µL of the sample was deposit for 5 min (15 µL during 15 min for AcS samples to compensate lower concentration) on a negatively charged grid (20 sec at 20 mA). The excess was wiped off. Negative staining was used to enhance contrast between the particles and the background (contaminants). The staining was performed using uranyl acetate. Samples preparation and imaging were performed with the help of Kelly Sears and Jeannie Mui from the Facility for Electron Microscopy Research (FEMR) of McGill University.

*Custom antibody microarray:* For the immuno-detection, a droplet of 400 pL of a solution of protein was deposited on an aldehyde glass slide (2-D Aldehyde slides, PolyAn GmbH, Berlin, Germany) by a non-contact piezo-microarrayer (Scienion) using a single nozzle (Type 1). A microarray 10 by 10 spots was printed, each slide had 16 replicates of each print. Samples were diluted to achieve a concentration of $1.5 \cdot 10^9$ EVs/mL (for cell supernatant samples), $10^{10}$ EVs/mL (for plasma samples) or for AcS plasma samples $10^8$ EVs/mL (AcS samples containing lower EV concentration) according to TRPS measurement. Detection was performed using a mixture of antibodies conjugated to biotin targeting CD63, CD9 and CD81, and streptavidin-AlexaFluor647. All prints were performed at room temperature and at 65% of humidity. The quantification of the fluorescent signal was performed using a microarray scanner (InnoScan 1100 AL Fluorescence scanner, Innopsys MBI Lab equipment). Detection signal was normalized by the buffer signal. If an image has a positive signal for a non-human IgG (named Neg in figures) the image is discarded.

*Analysis:* For plasma samples, the normalized final concentration of extraction is the final concentration after extraction related to initial and final volumes of samples:

$$[normalized\ final\ concentration\ of\ extraction] = \frac{Volume_{final}}{Volume_{initial\ of\ plasma}} [final\ concentration\ of\ the\ sample]$$

(eq1)

### Tables and Figures:

*Table S1 Volumes of the samples before and after extraction by UC, SEC, AcS and LoDF for samples of cell supernatant (CS) and plasma*

| Extraction method | UC | | SEC | | AcS | | LoDF | |
|---|---|---|---|---|---|---|---|---|
| Sample type | **CS** | **plasma** | **CS** | **plasma** | **CS** | **plasma** | **CS** | **plasma** |
| Initial volume | 2 mL | 1 mL | 2 mL | 500 µL | 2 mL | 80 µL | 2 mL | 100 µL |
| Final volume | 150 µL | 150 µL | 500 µL | 500 µL | 90 µL | 170 µL | 200 µL | 200 µL |

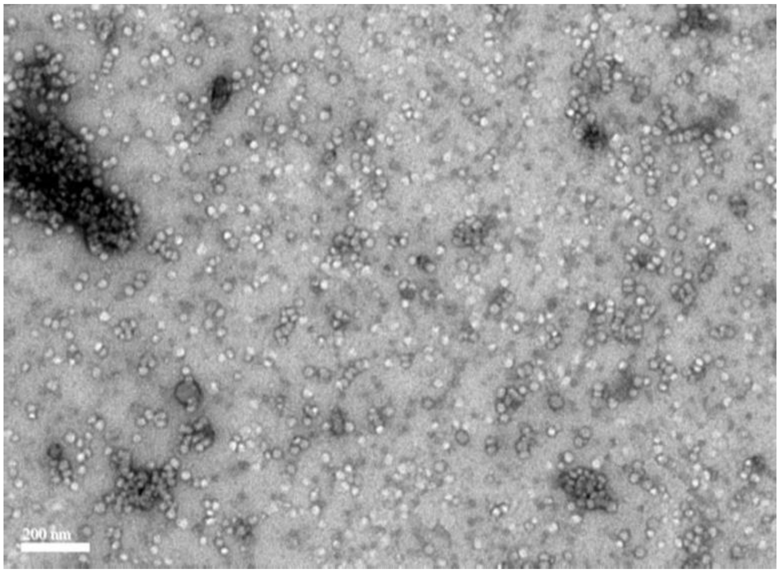


*Figure S1: TEM image of LoD extracted EVs from cell supernatant* (200nm scale bar)

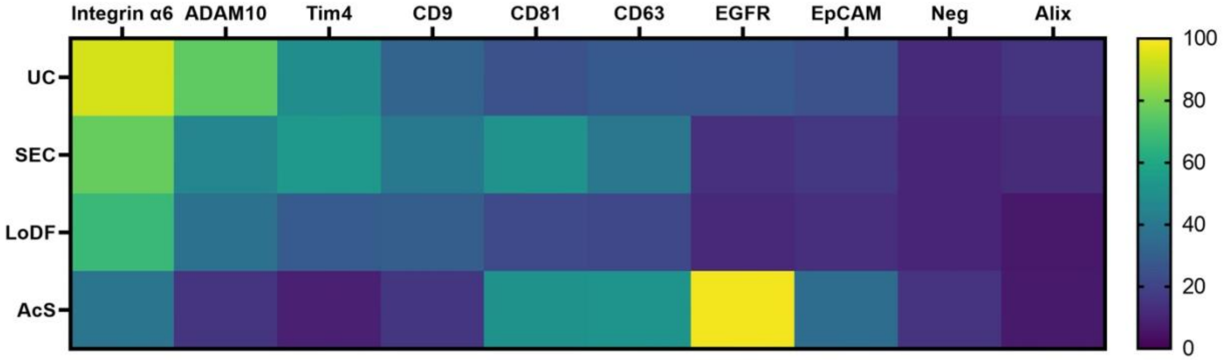


*Figure S2: Antibody microarray-based profiling of protein expression of EVs extracted from A431 CS. EVs extracted using each of the four methods were incubated on the microarray with the respective capture antibodies (top line), and then probed with a cocktail of fluorescently detection antibodies against CD9, CD63, CD81 (measured fluorescent signal). The signal is the normalized by the highest logarithmic value of the signal*

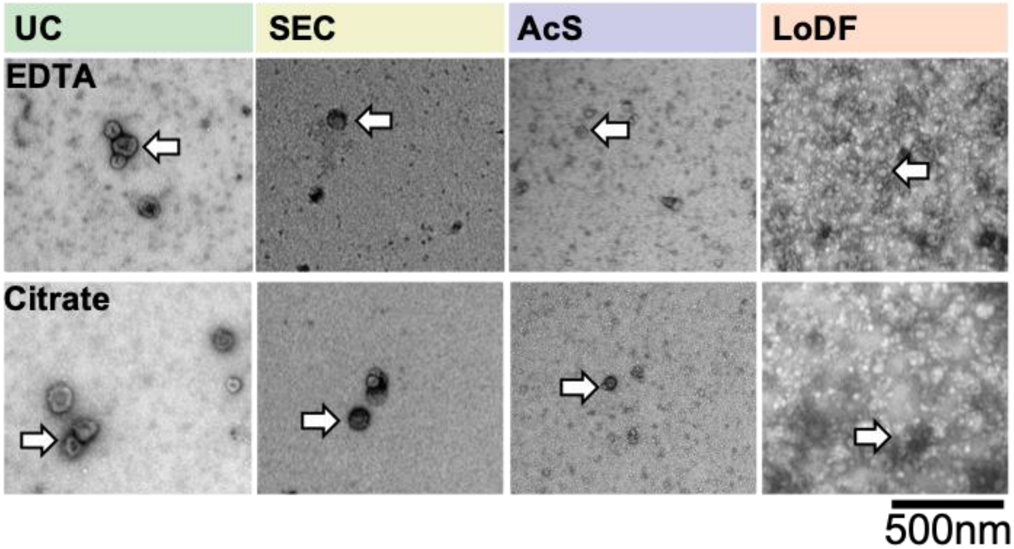


Figure S3. TEM images of EVs extracted from plasma samples processed using EDTA and Citrate tubes and extracted by UC, SEC, AcS and LoDF (SPP3 protocol). White arrows point to EVs. The images reveal the overall concentration of EVs, whether their shape has been preserved, and presence of contaminants. In all samples, the circularity and integrity of EVs was preserved. A higher concentration of particles, including non-EV particles, was observed in LoDF samples and a lower concentration in AcS samples. A high contamination by non-EV particles is noticeable for LoDF samples.

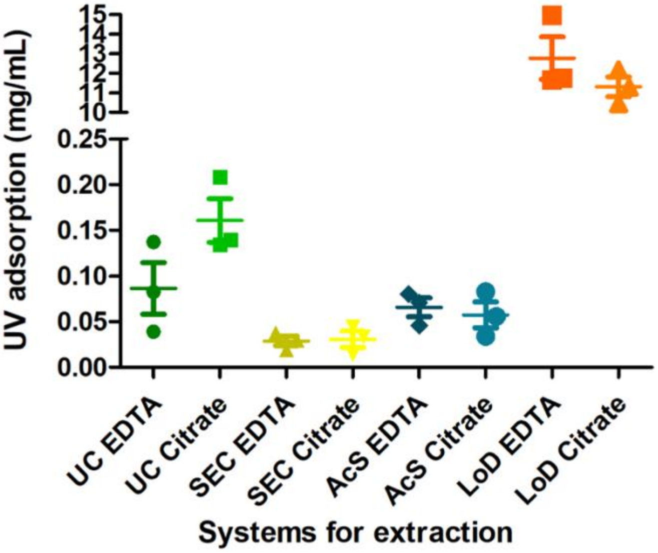


*Figure S4: Characterization by nanodrop of EVs extracted by UC, SEC, AcS and LoDF for EDTA and Citrate plasma sample*

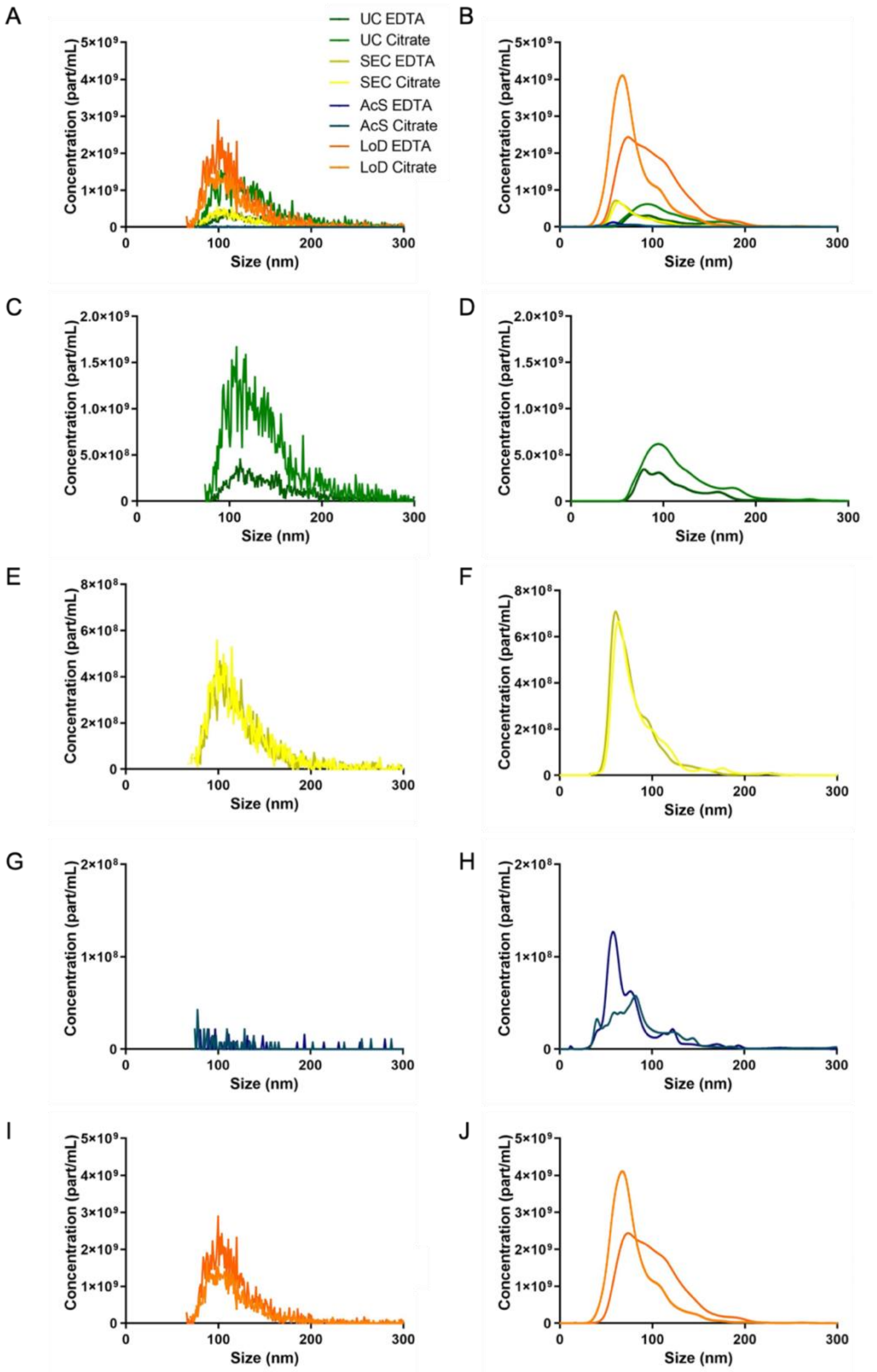


*Figure S5: Characterization of the size distribution of particles extracted by UC, SEC, AcS and LoDF measured by TRPS (A,C,E,G, I) and NTA (B, D, F, H, J) for EDTA and Citrate plasma sample.*

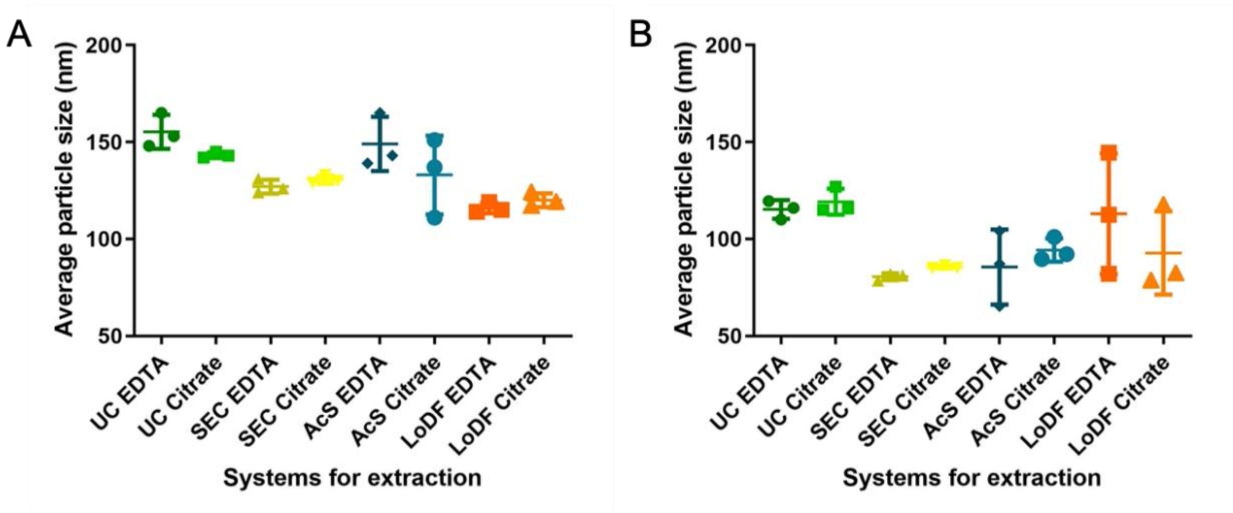


*Figure S6: Average size of the particles extracted by UC, SEC, AcS and LoDF measured by TRPS (A) and NTA (B) for EDTA and Citrate plasma sample.*

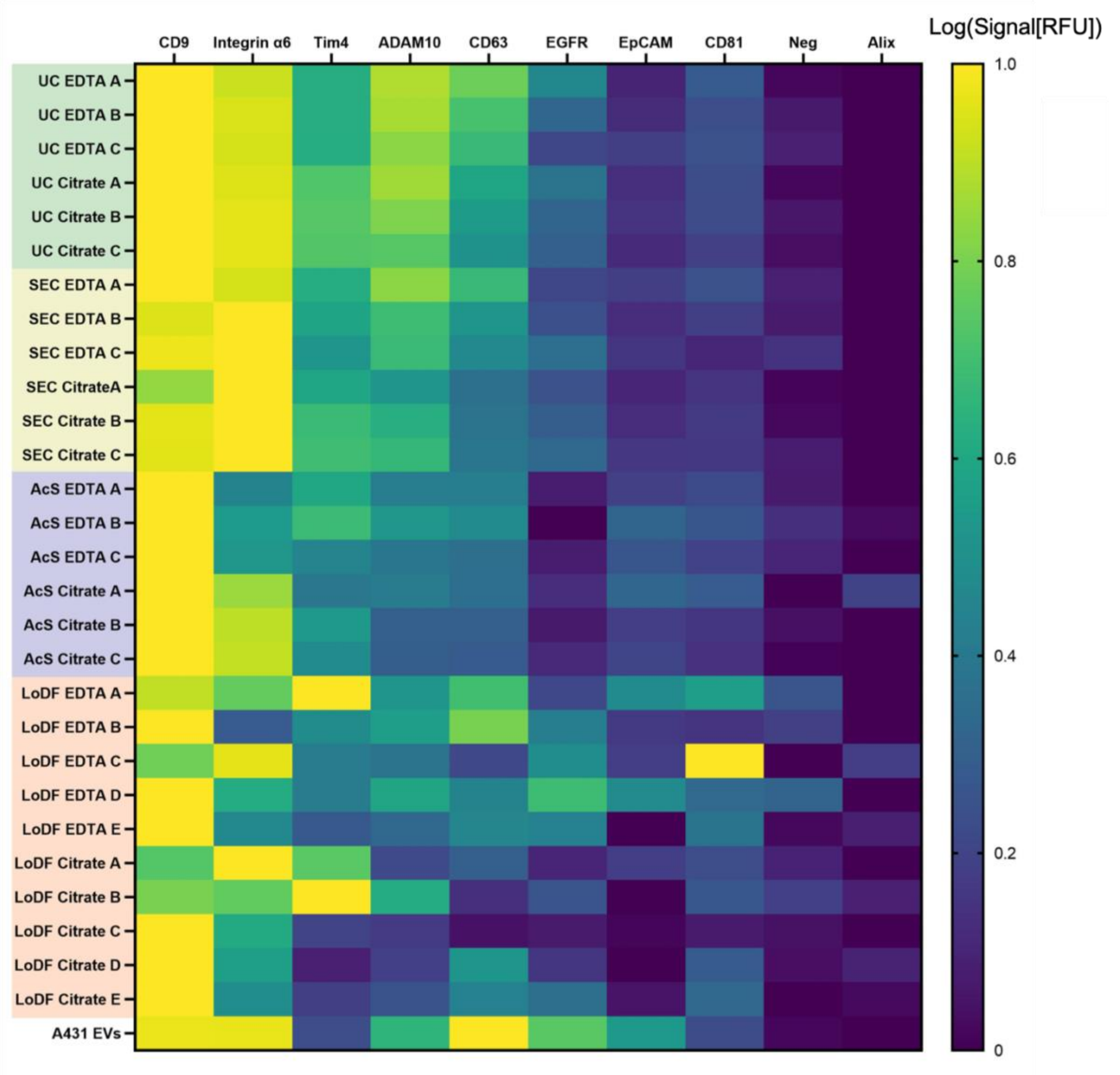


*Figure S7: Protein expression in EVs from A431 cell supernatant spiked into plasma profiled by antibody micro-array. EVs extracted using each of the four methods were incubated on the microarray with the respective capture antibodies (top line), and then probed with a cocktail of fluorescently detection antibodies against CD9, CD63, CD81 (measured fluorescent signal). The signal is the normalized by the highest logarithmic value of the signal*
